# FROM FOREST TO SAVANNA AND BACK TO FOREST: EVOLUTIONARY HISTORY OF THE GENUS *Dimorphandra* (LEGUMINOSAE)

**DOI:** 10.1101/2023.01.16.524261

**Authors:** Vinicius Delgado da Rocha, Thaís Carolina da Silva Dal’Sasso, Christina Cleo Vinson Williams, Marcelo Fragomeni Simon, Marcelo Leandro Bueno, Luiz Orlando de Oliveira

## Abstract

The tree genus *Dimorphandra* comprises 26 species, which are circumscribed into three subgenera. The subgenus *Dimorphandra* is associated with both rainforests (Amazon and Atlantic Forest) and savanna-like vegetation (Cerrado); whereas the subgenera *Pocillum* and *Phaneropsia* are restricted to the Amazon. We obtained DNA sequence data from six gene regions of the chloroplast genome (cpDNA) and the nuclear internal transcribed spacer (ITS) from 17 species of *Dimorphandra* and 12 closely related species. Bayesian phylogeny and haplotype network analyses together with both ancestral area reconstructions and ecological niche modeling allowed for exploring the late evolutionary history of the genus *Dimorphandra*. Species within the subgenus *Phaneropsia* were more closely related to species of the genus *Mora* than to the remaining congeners in the plastid tree (but not in the ITS tree), casting doubts on the monophyly of *Dimorphandra*. Such incongruence may be the result of incomplete lineage sorting of ancient polymorphisms. Amazonian lineages (subgenera *Pocillum* and *Phaneropsia*) were highly polymorphic and divergent; whereas lineages from either the Cerrado or the Atlantic Forest were genetically depauperate. The Amazon seems to be the likely source of the lineage that gave rise to the extant species of *Dimorphandra* of the Cerrado. In turn, a lineage that occupied the Cerrado likely gave rise to the extant species that occur in the Atlantic Forest. Habitat shifts may have been a key driving force that shaped the late evolutionary history of *Dimorphandra*.

## INTRODUCTION

In South America, the three main biogeographic domains (herein referred to as Amazon, Cerrado, and Atlantic Forest, respectively) occupy each a vast territory, comprise many ecosystems, and intermingle along their borders. The Amazon is home to the largest tropical forest in the world, which extends across vast areas in northern Brazil, the Guiana Shield, Colombia, Ecuador, Peru, and Bolivia (Cardoso et al. 2017). Along its southeastern border, the Amazon meets the Cerrado. The Cerrado runs as a diagonal domain across central Brazil and parts of Paraguay and Bolivia; it includes several types of vegetation, with the predominance of the savanna ecosystems (Pennington et al. 2000). In sequence, the Cerrado meets the Atlantic Forest, which extends along the eastern coast of Brazil and reaches eastern Paraguay and northeastern Argentina (Tabarelli et al. 2005). These three biogeographic domains share many phylogenetically related species of plants. The Cerrado shares flora with both the Amazon and the Atlantic Forest (Oliveira-Filho and Ratter 1995; Méio et al. 2003). Despite their geographic distance, the Amazon and Atlantic Forest domains share elements of their vegetation (Maciel et al. 2017; Terra-Araujo et al. 2015).

Geological and climatic events have shaped the landscapes of South America and influenced the diversification of biota over time (Pennington and Dick 2010). During the Oligocene-Pliocene periods, a sequence of geological events resulted in the Andean uplift, which in turn changed the hydrology, soil fertility, and climate of the Amazon and surrounding regions (Hoorn et al. 2010). Ancient episodes of climate changes together with vegetations shifts led to alternating cycles of isolation and reconnection among ancestral populations; intensely they triggered diversification events in both plants and animals within the Neotropics, including the Amazon (Leite and Rogers 2013), Cerrado (Bueno et al. 2017), and Atlantic Forest (Bacci et al. 2022). In the past, specifically from the Miocene to the Pleistocene, networks of forest and other type of corridor vegetation likely connected the Amazon to both the Cerrado and the Atlantic Forest (Oliveira-Filho and Ratter 1995; Ledo and Colli 2017; Machado et al. 2018). These connections allowed for intensive migrations of biota amongst these biogeographic domains (Buzatti et al. 2018; Fabrício Machado et al. 2021). Over time, many dispersal events occurred from the Amazon toward the Cerrado, and from the Amazon towards the Atlantic Forest; certainly, the Amazon was an important source of lineages to surrounding areas (Antonelli et al. 2018). On the other hand, the Amazon and the Atlantic Forest have both received lineages from the Cerrado (Aguiar et al. 2019; Gonçalves et al. 2020; Lima et al. 2021), but in a lower frequency. Dated molecular phylogenies and biogeographic analyses provided evidence that dispersal events were frequently associated with habitat shifts from forest and savanna or vice-versa. It seems that ecological transition between savanna and forest was a key-factor for the evolution of many plants and animals in the Neotropics (Simon and Pennington 2012; Terra-Araujo et al. 2015; Simon et al. 2016; Antonelli et al. 2018; Aguiar et al. 2019).

*Dimorphandra* Schott (Leguminosae) is a tree genus distributed across different habitats in the Neotropics, such as rainforests (Amazon and Atlantic forests) and savannas (Cerrado). The genus was first described in 1827, having *D. exaltata* Schott as the type species (Silva 1986). Species of *Dimorphandra* share morphological traits, such as bipinnate leaves, paniculate inflorescences that are erect, hermaphrodite flowers with a narrow receptacle, presence of five stamens and five staminodes, and corollas that consist of five petals (Silva 1986; Silva 2019). Currently, *Dimorphandra* comprises 26 described species distributed into three subgenera, which are well defined on morphological grounds (Silva 1986; Silva 2019). The fruit morphology is the most striking feature that distinguishes the subgenera. Morphologically, fruits traits for each subgenus are as following: a) subgenus *Pocillum* Tul.: woody fruits, flattened, falcate, dehiscent, ventral margin is large, and dorsal margin is thin; b) subgenus *Phaneropsia* Arch.: woody fruits, linear or falcate, and dehiscent; c) the subgenus *Dimorphandra:* elongated fruit with a large thickness, linear or slightly curved, and indehiscent (Silva 1986).

With ten species, the subgenus *Pocillum* is restricted to the Amazon; mostly on the northern side of the Amazon River, across the Guiana Shield. The subgenus *Phaneropsia*, with 5 species, is distributed from northern to western South America; species distribution occurs primarily across the Amazon. Finally, the subgenus *Dimorphandra*, with 11 species, exhibits widespread distribution, from northern South America to southeastern Brazil, reaching both the Cerrado and the Atlantic Forest (Silva 1986). Although the geographic distribution of the genus has not been thoroughly characterized, it is already clear that *Dimorphandra* shows uneven distribution across each of the three main biogeographic domains: Amazon, Cerrado, and Atlantic Forest.

In the Amazon, some congeners show widespread occurrence, such as *D. pennigera* and *D. macrostachya*. Meanwhile, many other species have been suggested to be narrow endemics, such as *D. davisii, D. urubuensis*, and *D. campinarum* (Silva 1986; Silva and Hopkins 2018). Indeed, there are several species (*D. davisii*, *D. dissimilis*, *D. gigantea*, *D. multifora*, *D. mediocris*, *D. ignea*, *D. loretensis*, *D.pullei*, *D. urubuensis*, and *D. williamii*) for which a handful of records, or even a single herbarium specimen, are available. Such low occurrence of herbarium records, which is related to the enormous geographic size of that biogeographic domain and overall low sampling effort, complicates the inference of whether a given species is actually a rare/endemic species or its presumed endemism is the result of undersampling. The accumulation of *Dimorphandra* species across the Amazon suggests that the genus probably originated in that region, and later expanded its geographic range to other biogeographic domains.

Two species commonly found across the Cerrado, *D. mollis* and *D. gardneriana*, are likely the two most widespread species within the genus (Silva 1986). Both *D. mollis* and *D. gardneriana* have attracted commercial interest because those species are sources of active chemical compounds. The flavonoid rutin plays a role as an antioxidant and an anti-inflammatory agent (Silva 1986; Cunha et al. 2009). The geographically narrow endemic species, *D. wilsonii* is considered to be seriously threatened with extinction; the species is found at some few neighboring sites of a Cerrado/Atlantic Forest ecotone in the central parts of Minas Gerais state (Fernandes and Rego 2014; Vinson et al. 2015).

Currently, *D. jorgei* and *D. exaltata* (the type species) are the only species known to occur within the Atlantic Forest, both of which detain restricted geographic distribution (Silva 1986). The evolutionary relationship among species of *Dimorphandra* from the Amazon and extra-Amazon species of *Dimorphandra* remains to be explored. *Dimorphandra* species of these two later domains share remarkable morphological similarities (Silva 1986). A recent study proposed that *D. wilsonii*, a species from Cerrado/Atlantic Forest ecotone is an interspecific hybrid between *D. mollis* from the Cerrado and *D. exaltata* from Atlantic Forest; *D. wilsonii* exhibited microsatellite alleles found in either *D. mollis* or *D. exaltata* (Muniz et al. 2020).

Considering its widespread occurrence in different vegetation domains, and its supposedly intricate speciation patterns influenced by ecological transitions together with both past climate changes and hybridization in ecotonal zones, *Dimorphandra* stands as a promising study group to investigate the diversification of the rich Neotropical tree flora. In the present investigation, we used an array of molecular phylogenetic analyses together with both ancestral area reconstructions and ecological niche modeling to shed light on the evolutionary history of the genus *Dimorphandra*. We began our study by questioning the monophyly of *Dimorphandra* as currently described and the genealogical relationships amongst members of *Dimorphandra* and amongst *Dimorphandra* and closely related genera. Next, we searched for evidence about which of the three main biogeographic domains of South America (Amazon, Cerrado, or Atlantic Forest) could be the most likely geographic source of the extant lineages of *Dimorphandra*; we also inquired whether habitat shifts may have contributed to the diversification of *Dimorphandra*. Subsequently, we used species distribution modelling to explore how climatic conditions may have shaped the geographic distribution of *Dimorphandra* during the last 130,000 years. The paleodistributional predictions may provide clues about expansion, reduction, and stability of potentially favourable conditions compared to present-day distribution of *Dimorphandra*. Finally, we investigated whether the narrow endemic congeners that occur in forested areas along the Atlantic coast of Brazil could be evolutionary relicts or, alternatively, could be recently derived species. The understanding of the evolutionary history of *Dimorphandra* will shed light on the origin and assembly of the tree flora in the diverse biogeographic domains of South America.

## MATERIALS AND METHODS

### Taxon sampling

Our sampling strategy covered most of the range of the genus *Dimorphandra*, and included sampling sites within three biogeographic domains (Amazon, Cerrado, and Atlantic Forest), including representatives of all three subgenera of *Dimorphandra*. We also sampled *D. wilsonii*, a species that is typically found within the Cerrado/Atlantic Forest ecotone.

A total of 59 specimens, representing 17 species of *Dimorphandra*, were represented in our study, including seven species of the subgenus *Dimorphandra* (*D. caudata*, *D. exaltata*, *D.gardneriana*, *D. jorgei*, *D. mollis*, *D. parviflora*, and *D. wilsonii*); three species of subgenus *Phaneropsia* (*D. conjugata*, *D. dissimilis*, and *D. unijuga*); and seven species of subgenus *Pocillum* (*D. campinarum*,*D. coccinea*, *D. cuprea*, *D. macrostachya*, *D. pennigera*, *D. polyandra*, and *D. vernicosa*) (Supplementary Table S1).

Additionally, we also included 12 species of nine closely related genera based on previous phylogenetic analyses (Bruneau et al. 2008): *Campisandra* (one species), *Dinizia* (one), *Entada* (one), *Mora* (two), *Peltophorum* (one), *Parkia* (two), *Schizolobium* (one), *Stryphnodendron* (one), and *Tachigali* (two) (Supplementary Table S1).

### DNA extraction, amplification, and sequencing

The DNA extraction and the polymerase chain reactions (PCRs) were carried out at the Laboratory of Molecular Biology and Phylogeography, Institute Biotechnology Applied to Agriculture (BIAGRO), Universidade Federal de Viçosa (UFV), Viçosa, Minas Gerais, Brazil. The total genomic DNA was extracted from either silica gel-dried leaves or herbarium specimens. In both cases, we used a previously described CTAB protocol (Riahi et al. 2010). Independent PCRs amplified six regions. There were five regions of the chloroplast genome (cpDNA) and a region of the nuclear genome. The five cpDNA regions were: the trnL-UAA intron (Taberlet et al. 1991), three intergenic spacers [*trnL* – *trnF* (Taberlet et al. 1991); *trnH* – *psbA* (Shaw et al. 2005); *psbB* – *psbF* (Hamilton 1999)], and a region of the gene Maturase K - *mat*K (Kuzmina et al. 2012). The nuclear region comprised the entire internal transcribed spacer (ITS) of the 18S-26S ribosomal RNA genes, including the 5.8S gene (White et al. 1990; Baum et al. 1998). The primers used during PCR were as listed (Supplementary Table S2).

For each of the five cpDNA regions, PCRs were carried out in a final volume of 25 μL, containing: Green 1X Go Taq (Promega) buffer; 17.5 μg/mL of BSA (*Bovine Serum Albumin);* 0.2 mM of each dNTP; 2 mM of MgCl2; 0.5 μM of each primer, 1.25 U of Go Taq Flexi (Promega), and approximately 50 ng of genomic DNA. The *psb*B – *psb*F region was amplified as follows: 95°C for 5 min (initial denaturation), followed 35 cycles of 95°C for 1 min, 58°C for 1 min (primer annealing) and 72° C for 90s, and final extension of 72°C for 5 min. The remaining four cpDNA regions were amplified as follows: 95 °C for 3 min (initial denaturation), followed 40 cycles of 95 °C for 1 min, 54 °C for 1 min (primer annealing), 72 °C for 1 min, and final extension of 72°C for 10 min.

The amplification of the ITS region was carried out in a final volume of 20 μL, with Green 1X Go Taq (Promega) buffer, 1.75 μL of DMSO (dimethyl sulfoxide), 0.25 mM of each dNTP, 2.5 mM of MgCl2, 0.62 μM of each primer, 1.25 U of Go Taq Flexi (Promega), and approximately 100 ng of genomic DNA. The amplification program consisted of 94 °C for 4 min (initial denaturation), followed 35 cycles of 94 °C for 1 min, 49-58°C for 1 min (primer annealing), 72 °C for 45 s, and final extension of 72 °C for 5 min.

Prior to sequencing, PCR products were purified using ExoSAP IT (USB; 3 μL of enzyme per 9 μL reaction). Sequencing was carried out at Macrogen Inc. in South Korea (www.macrogen.com).

### Assembly of datasets

Sequence chromatograms were visually inspected, and the sequences were aligned using the software Sequencer v. 4.8 (Gene Code Corp), with manual edition of gaps for the presence of regions of insertions or deletions (indel). In the chromatograms, sites with double peaks were coded according to the nucleotide ambiguity codes, as defined by IUPAC (International Union of Pure and Applied Chemistry). Sequences were deposited in the GenBank (Supplementary Table S3).

For phylogenetic analyses, we assembled two datasets. Dataset A (N=73; 2622 bp) contained the sequences of the five cpDNA regions concatenated (trnL-UAA, 495 bp; *trn*L – *trn*F, 425 bp; *psb*B – *psb*F, 560 bp, *trn*H – *psb*A, 481 bp; and *mat*K, 661 bp). Dataset B (N = 53; 638 bp) contained sequences of ITS region.

For network analyses, the following source of information was removed from the datasets: (1) ambiguous sites; (2) sites that displayed more than two character state, because they violated the infinite-sites model (Kimura 1969) and could introduce homoplasy in the network; and (3) indels that consisted of more than four mononucleotide repeats (AAAA, for instance). Regardless of the length, each indel was coded as a fifth character state to become a single mutation. Finally, we obtained two additional datasets. Dataset C (N = 55; 1782 bp) contained the concatenated sequences of cpDNA regions (trnL-UAA, 448 bp; *trn*L – *trn*F, 367 bp; *psb*B – *psb*F, 549 bp; *trn*H – *psb*A, 418 bp). Dataset D (N = 56; 446 bp) harbored the sequences of ITS region, including twelve additional sequences of *D. mollis* that were available from the GenBank (accessions KT970387–KT970398).

### Phylogenetic analyses

To uncover the phylogenetic relationships among species of *Dimorphandra*, we used two methods: Bayesian inference and maximum likelihood (ML).

Prior to the Bayesian phylogeny, we selected the best-fit evolutionary model for each gene region, independently, using the Akaike Information Criterion (Akaike 1998), as implemented in the software MrModeltest v2.3 (Nylander 2004). The best-fit models for each region were the following: GTR+G (trnL-UAA, *trn*H – *psb*A, and *mat*K), GTR+I+G (*trn*L – *trn*F and ITS), and GTR+I (*psb*B – *psb*F). Independent Bayesian phylogenies were performed for dataset A and dataset B, using the software MrBayes v3.1.2 (Ronquist and Huelsenbeck 2003). In MrBayes, each analysis consisted of two simultaneous runs of 10 million generations; one cold and seven heated chains in each run; the temperature parameter was of 0.23. Sampling was performed every 10,000 generations; the first 250 trees were discarded as burn-in. The average standard deviation of split frequencies at end of each run was less than 0.01. We examined the effective sample size (ESS) values for all statistics, using software Tracer v.1.5 (Rambaut and Drummond 2009). ESS values were above 200 and indicated convergence of the Markov chains. Finally, we obtained a 50% majority-rule consensus for each dataset (A and B).

For the ML analyses, we also inferred the best-fit evolutionary for each gene region, separately, using Bayesian Information Criterion (BIC) from the ModelFinder 2 (Kalyaanamoorthy et al. 2017), as implemented in the software IQ-TREE (Nguyen et al. 2015). The best-fit evolutionary models for each gene region were as following: TVM+F+G4 (trnL-UAA and *mat*K), TVM+F+R2 (*trn*L – *trn*F), TVM+F+I (*psb*B – *psb*F), GTR+F+G4 (*trn*H – *psb*A), and TIM3+F+I+G4 (ITS). Using the software IQ-TREE (Nguyen et al. 2015), the ML analyses were performed independently for dataset A and dataset B. Each ML analysis was performed using independent runs and 1000 ultrafast bootstrap replicates, having 0.99 as the minimum correlation coefficient for the bootstrap convergence criterion (Hoang et al. 2018).

Finally, we used FigTree v1.2.4 (Rambaut 2009) to visualize the phylogenetic trees, which were obtained from both Bayesian and ML analyses.

### Genealogical network

Both dataset C and dataset D allowed us to infer the genealogy relationships among lineages of *Dimorphandra*. We reconstructed haplotype networks using the Median Joining network method (Bandelt et al. 1999) as implemented in the software Network v.5.0.03 (Fluxus Technology Ltd).

### Ancestral area reconstruction

The ancestral area reconstruction was carried out using the Bayesian Binary MCMC (BBM) analysis as implement in software RASP v.4.2 (Yu et al. 2015). The geographic areas were defined on the basis of the current distribution range of the *Dimorphandra* species, using information from Global Biodiversity Information Facility (Gbif.org 2018) and Flora do Brasil 2020 (Souza and Lima 2020). Based on the biogeographic domains, we defined four areas: Amazon, Cerrado, Cerrado/Atlantic Forest ecotone, and Atlantic Forest.

For the BBM analysis, we used as input a set of 1000 output trees and a consensus tree obtained in Bayesian phylogeny with cpDNA data (dataset A). The BBM analysis was performed using model F81 + G; two runs of 6 million generations each; ten chains, and temperature parameter of 0.2. Sampling every 6.000 generations, and the 250 first samples were discarded as burn-in. The maximum number of areas for each node was set to three.

### Ecological Niche Modeling

From public repositories, we obtained georeferenced occurrences for seven species of *Dimorphandra* (*D. exaltata, D. gardneriana, D. jorgei, D. macrostachya, D. mollis, D. pennigera*, and *D. wilsonii*). The georeferenced occurrences were carefully chosen to cover the range of each species. The number of georeferenced occurrences that was collected per species varied from 13 to 50, and depended on the number of specimens that were available within the public repositories and how carefully the deposit was originally curated (Supplementary Table S4).

To analyze the ecological Niche Model (ENMs) of the seven species of *Dimorphandra*, used the 19 climate variables extracted from WorldClim database (Hijmans et al. 2005) for all geographical coordinates at a spatial resolution of 1 km, that reflect various aspects of temperature, precipitation, and seasonality, which are likely to be important in determining species distributions. We used a stepwise procedure implemented in the R usdm package (Naimi and Araújo 2016) in R 3.6.3 (R Development Core Team 2020) to test the issue of multicollinearity among the environmental variables by estimating the variance inflation factor (VIF) and retained only the variables with VIF < 10 (Graham 2003). This reduced our number of environmental predictors to eight (Supplementary Table S5).

To infer the palaeodistribution of late Quaternary climatic change of the seven species of *Dimorphandra* distribution, we produced projections of the suitability of occurrence during the Present (0 ka pre-industrial), Last Glacial Maximum (LGM – 21 ka BP), and Last Interglacial (LIG – 130 ka BP) time periods based on climatic simulations (Hijmans et al. 2005). For the Last Glacial Maximum (21 ka, LGM), we employed the Community Climate System Model – CCSM4 (Gent et al. 2011) and MIROC-ESM (Watanabe et al. 2011), which represent downscaled climate data from simulations with Global Climate Models (GCMs) based on the Coupled Model Intercomparison Project Phase 5 (CMIP5; Taylor et al. 2012). We summed the projections of the species for each time period, which together represent the probability of occurrence during that time period. The paleo-climatic model for the Last Interglacial (120 ka, LIG) used the approach of Otto-Bliesner et al. (2006).

We fitted ENMs for each species using four modelling algorithms implemented in the sdm package in R (Naimi and Araújo 2016). These were maximum entropy (MaxEnt) (Phillips et al. 2006); random forests (rf) (Breiman 2001); generalized linear models (glm) (McCullagh and Nelder 1989) and generalized additive models (gam) (Yee and Mitchell 1991). These methods were used to link the current environmental conditions to species presence and absence data, and subsequently, to predict and map the spatial distribution of the species for the current and paleoclimatic projections. All models were calibrated using presence data, which were combined with 10,000 randomly selected pseudo-absence records for each species across the study area, generated with the R sdm package (Naimi and Araújo 2016). Rather than relying on individual modelling algorithm approaches, we built ensemble models combining multiple replicates of several different modelling algorithms to represent alternate possible states of the system being modelled (Araújo and New 2007). Due to their combined power, ensemble models are widely accepted to provide more accurate results than single models (Forester et al. 2013).

To assess the predictive capacity of the models, we divided the data for each species into a training set (75% of occurrences) and a test or validation set (25% of occurrence) performed with ten replicates subsampling method. To validate the produced models, we used the Area Under Curve (AUC) as a threshold-independent measure and the True Skill Statistic (TSS) as threshold-dependent accuracy measures (Allouche et al. 2006; Liu et al. 2009), as well as the correlation coefficient (COR) (Bradley 1997)

## RESULTS

### Phylogenetic inferences

After exclusion of ambiguous sites, the five concatenated cpDNA regions comprised a total of 2615 bp; meanwhile the ITS region contained 600 bp. Overall, the five cpDNA regions and the ITS region provided informative polymorphisms, thus they were useful for phylogenetic analyses at the species level. The number of segregating sites varied from 38 (cpDNA dataset) to 63 (ITS dataset). Among the species of *Dimorphandra*, the average number of nucleotide differences varied from 3.63 (cpDNA) to 10.84 (ITS). An overview of the measures of nucleotide diversity was presented for both cpDNA and ITS datasets (Table 1).

**Table 1.** Measures of nucleotide diversity for the cpDNA dataset and ITS dataset among 17 species of *Dimorphandra*. The cpDNA dataset was obtained from the concatenation of five regions: an (trnL-UAA) intron, three intergenic spacers (*trn*L–*trn*F, *trn*H–*psb*A, and *psb*B–*psb*F) and a region of the Maturase K gene.

|  | **cpDNA** | **ITS** |
| --- | --- | --- |
| Number of sequences | 73 | 53 |
| Number of total base pairs | 2615 | 600 |
| Number of segregating sites | 38 | 63 |
| Number of indels | 28 | 26 |
| Number of haplotypes | 19 | 19 |
| Haplotype diversity | 0.841 | 0.880 |
| Nucleotide diversity | 0.006 | 0.036 |
| Average number of nucleotide differences | 3.63 | 10.84 |

Using Bayesian inference and ML analyses, we reconstructed the phylogenetic relationships among 17 species of *Dimorphandra* and 12 species of nine closely related genera, with *Entada polystachya* as outgroup. There were two independent phylogenetic reconstructions, based on the cpDNA dataset and ITS dataset, respectively. The Bayesian phylogenies (Figs. 1 and 2) and ML phylogenies (Supplementary Figs. S1 and S2) exhibited congruent topologies, showing high support for most of the internal nodes. All major clades were recovered in both Bayesian and ML analyses. Because of this congruence, we used only the Bayesian phylogenies during subsequent discussions.

**Fig.1.**
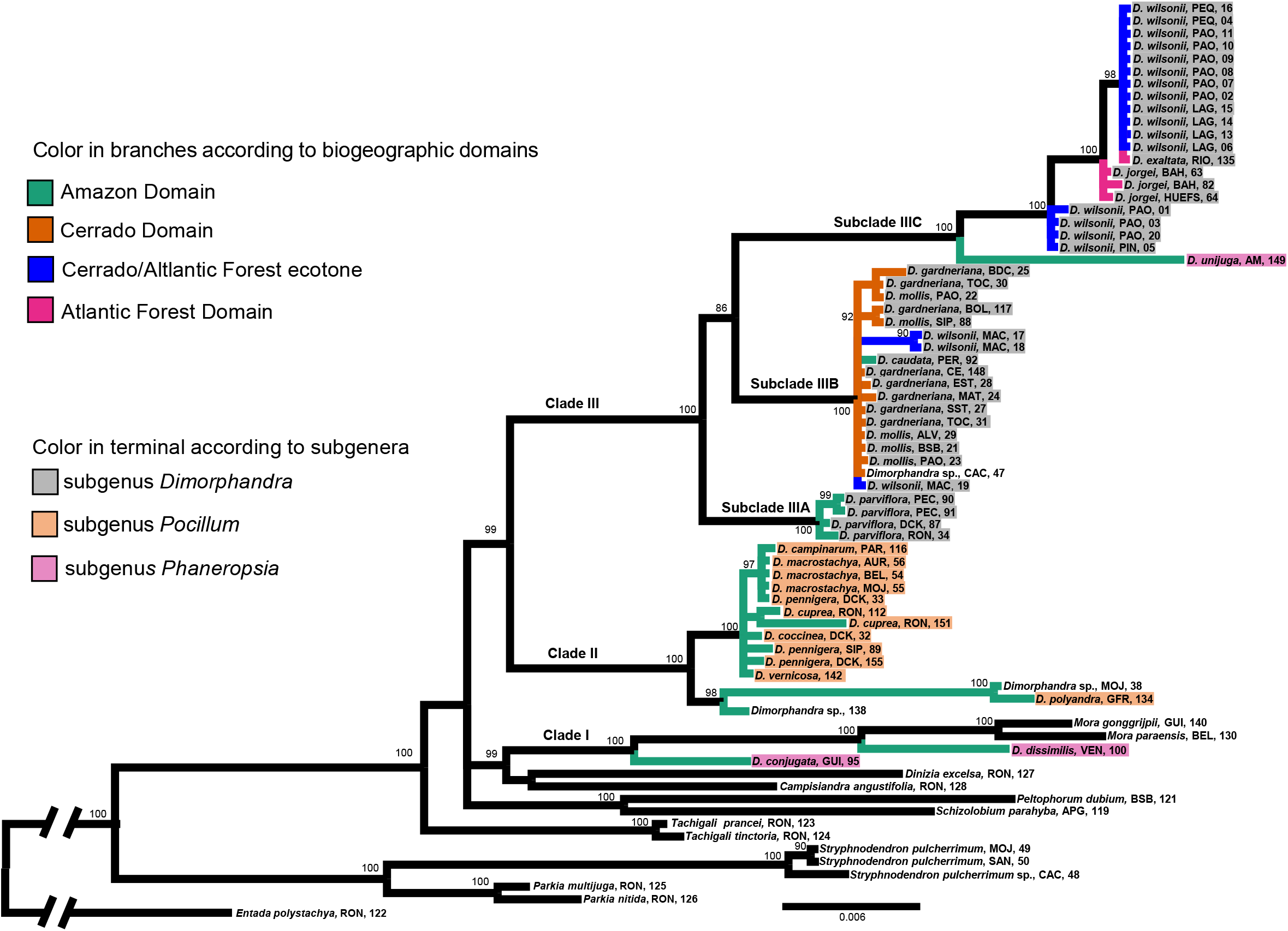
Bayesian phylogeny (consensus tree) showing relationship among 17 species of *Dimorphandra* (59 specimens) and nine related genera (14 specimens). The dataset (2615 pb) was obtained from the concatenation of five cpDNA regions: an intron (trnL-UAA), three intergenic spacers (*trn*L–*trn*F; *trn*H–*psb*A and *psb*B–*psb*F) and a region of the gene Maturase K. Branch lengths are drawn to scale. Nodal support values are given as posterior probabilities (%) above the branches (when ≥ 85). Scale bar corresponds to the expected number of substitutions per site.

**Fig.2.**
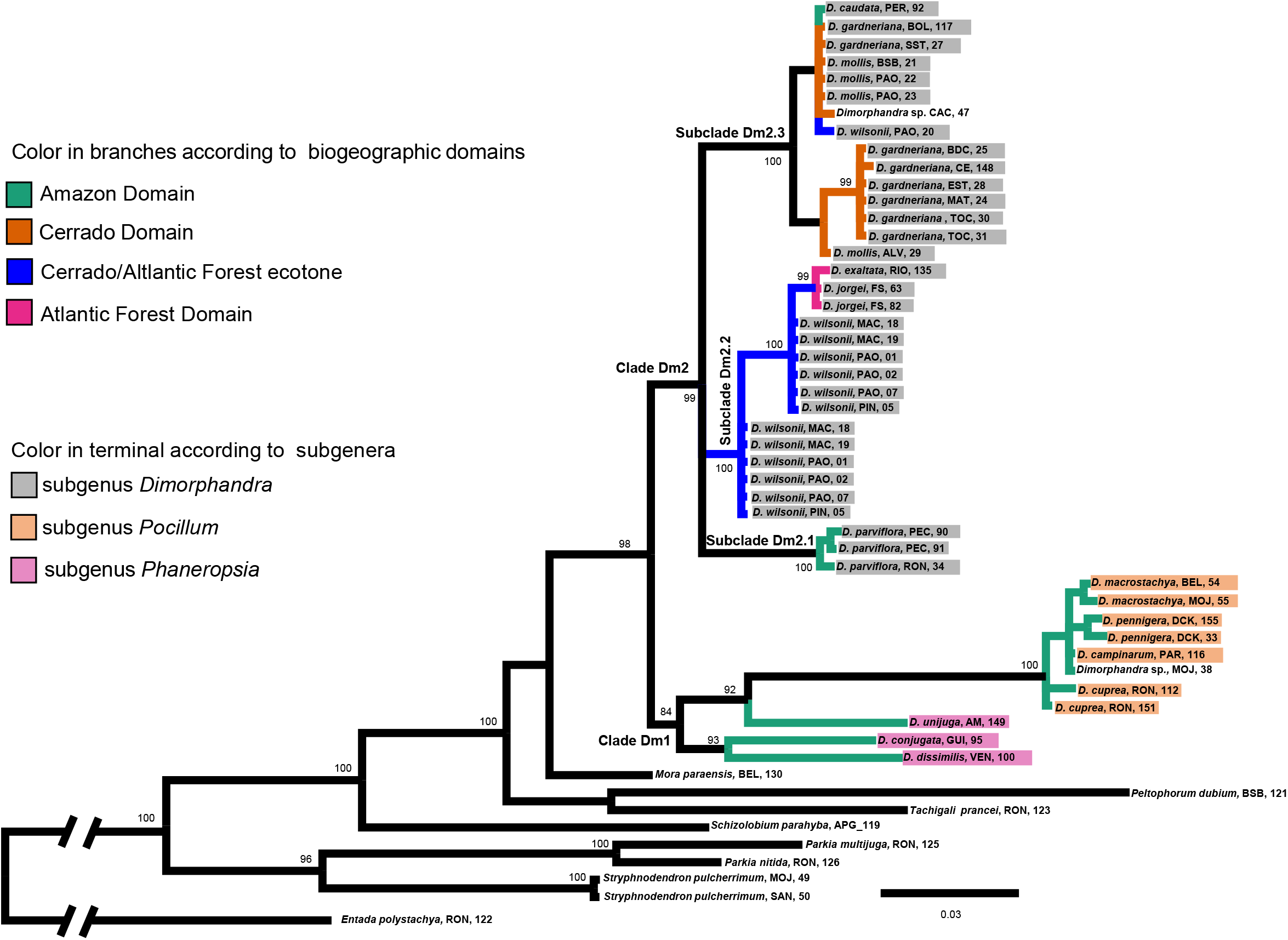
Bayesian phylogeny (consensus tree) showing relationship among 14 species of *Dimorphandra* (44 specimens) and seven related genera (9 specimens). The dataset (638 pb) was resulted from nuclear region (ITS). Branch lengths are drawn to scale. Nodal support values are given as posterior probabilities (%) above the branches (when ≥ 85). Scale bar corresponds to the expected number of substitutions per site.

The Bayesian consensus tree based on cpDNA dataset (Fig. 1) split the 17 species of *Dimorphandra* into three main clades, each of which was highly supported, with posterior probability (PP) = 100%. Herein, we will refer to these three major clades as Clades I, II and III, respectively. Clade I included two species of subgenus *Phaneropsia* (*D. conjugata* and *D. dissimilis*) plus *Mora paraensis* and *M.gonggrijpii*, all of which were from the Amazon, rendering *Dimorphandra* non-monophyletic. Clade II consisted exclusively of species that belong to subgenus *Pocillum* (*D. campinarum*, *D. coccinea*, *D. cuprea*, *D. macrostachya*, *D. pennigera*, *D. polyandra*, and *D. vernicosa*); all of which were from the Amazon. Clade III, which showed a sister relationship to Clade II, split further into three subclades. Subclade IIIA showed a sister relationship to both Subclades IIIB and IIIC, each of which having maximum node support values (PP=100%). Subclade IIIA contained the Amazonian *D. parviflora* as its single species member. Subclade IIIB grouped one species from the Amazon (*D. caudata*), one species from the Cerrado/Atlantic Forest ecotone (*D. wilsonii*) and two species from the Cerrado (*D. mollis* and *D. gardneriana*). At the most derived placement, Subclade IIIC brought together species either from subgenus *Dimorphandra* or subgenus *Phaneropsia* occurring in three domains: Amazon (*D. unijuga*), Cerrado/Atlantic Forest ecotone (*D. wilsonii*), and Atlantic Forest (*D. jorgei* and *D. exaltata*).

Meanwhile, the Bayesian consensus tree obtained from the ITS dataset (Fig. 2) recovered *Dimorphandra* as monophyletic (PP=98%) and split it into two major clades: Dm1 (PP=84%) and Dm2 (PP=99%), respectively. Clade Dm1 brought together seven species from Amazon: three species were from subgenus *Phaneropsia* (*D. dissimilis*, *D. conjugata*, and *D. unijuga*) and four species were from subgenus *Poccillum* (*D. cuprea*, *D. campinarum*, *D. pennigera*, and *D. macrostachya*). Members of Clade Dm2 belong to subgenus *Dimorphandra* exclusively. With strongly supported nodes (PP=100%), Subclade Dm2 split further into three subclades (Dm2.1, Dm2.2, and Dm2.3, respectively). Subclade Dm2.1 contained *D. parviflora*, a species from the Amazon, as its single member. Meanwhile, subclade Dm2.2 contained a species from the Cerrado/Atlantic Forest ecotone (*D. wilsonii*) together with two species from Atlantic Forest (*D. jorgei* and *D. exaltata*). Within subclade Dm2.2, *D. jorgei* and *D. exaltata* formed a terminal clade apart from *D. wilsonii*. Subclade Dm2.3 included two species from the Cerrado (*D. mollis* and *D. gardeneriana*), one species from Amazon (*D. caudata*), and one species from the Cerrado/Atlantic Forest ecotone (*D. wilsonii*).

### Haplotype networks

The haplotype network based on the cpDNA dataset depicted the genealogical relationships among 19 haplotypes, which were recovered from 55 specimens of *Dimorphandra* (Fig. 3). With a single exception (haplotype 19; see below), the placement of nearest haplotypes was arranged according to the three subgenera: *Pocillum* (haplotypes 1-9), *Phaneropsia* (haplotypes 10 and 11), and *Dimorphandra* (haplotypes 12-18). Moreover, the haplotype network was geographically structured to a certain degree, as the placements of nearest haplotypes follow closely the biogeographic domains in which species were collected.

**Fig.3.**
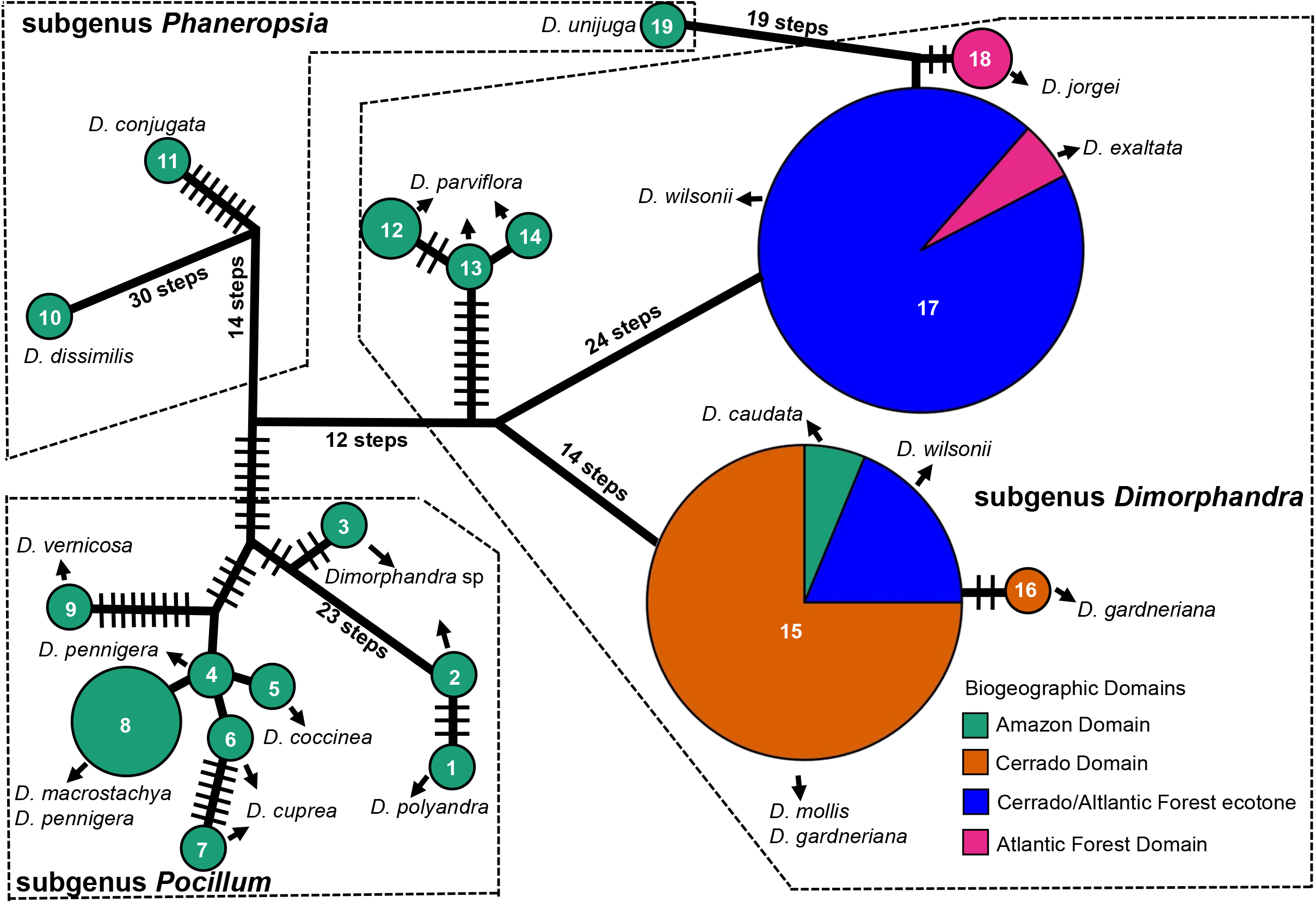
Median-joining network for 55 specimens of *Dimorphandra*. The dataset (1782 pb) was obtained from the concatenation of four cpDNA regions: an (trnL-UAA) intron, three intergenic spacers (*trn*L–*trn*F, *trn*H–*psb*A, and *psb*B–*psb*F). Each circle represents a given haplotype (coded with number); size is proportional to relative frequencies. Numbers of mutational steps are indicated with bars when there is more than one (unless indicated otherwise).

Over the extension of the network topology, species typically found in the Amazon (coded green) were genealogically closely related (see haplotypes 1-14). A general trend of the Amazonian species was the large number of mutation steps that separated nearby haplotypes (up to 30 steps), an indication that the cpDNA genome of these species detain a large number of autapomorphies. The placement of haplotype 19 contrasted sharply with the placement of other haplotypes that were recovered from Amazonian species. Haplotype 19 was uncovered from *D*. *unijuga*, an Amazonian species of the subgenus *Phaneropsia*; however this haplotype is genealogically more closely related to haplotypes found in the species of the Cerrado/Atlantic Forest ecotone (20 mutational steps were required to link to derived haplotype 19 to the central haplotype 17).The large number of haplotypes and the large number of mutational steps that were necessary to connect the haplotypes were features that characterized the network component that was obtained from the Amazonian species of *Dimorphandra*. These features stand in sharp contrast to those of the species of the remaining domains.

In both Cerrado and Atlantic Forest, and in their ecotone, the general trend was the presence of two haplogroups, each of which centered around a haplotype (15 or 17). A total of 38 mutational steps separated haplotype 15 from haplotype 17. Within each of these two haplogroups, few mutation steps separated nearby haplotypes (up to three steps). The only exception was haplotype 19, which required 20 steps to connect to the central haplotype 17. Two species of the Atlantic Forest (*D. exaltata* and *D. jorgei*) exhibited haplotypes (17 and 18) that were closely related to the haplotypes found in the ecotone. Interestingly, *D. wilsonii*, a species of the ecotone, contained two distantly related haplotypes (15 and 17). In addition to the species found in the Cerrado, the Cerrado/Atlantic Forest ecotone, and the Atlantic Forest, the each of these haplogroups contained a species from the Amazon (*D. unijuga* and *D. caudata*, respectively).

The haplotype network that was based on the nuclear gene (ITS) depicted the genealogical relationships among 18 haplotypes, which were recovered from 56 specimens of *Dimorphandra* (Fig. 4). To a certain degree, the distribution of haplotypes in the ITS-based network also was associated with subgenera: *Pocillum*, (haplotypes A-E), *Phaneropsia* (H-G), and *Dimorphandra* (I-R) and also associated with the biogeographic domain in which the species occurred.

**Fig.4.**
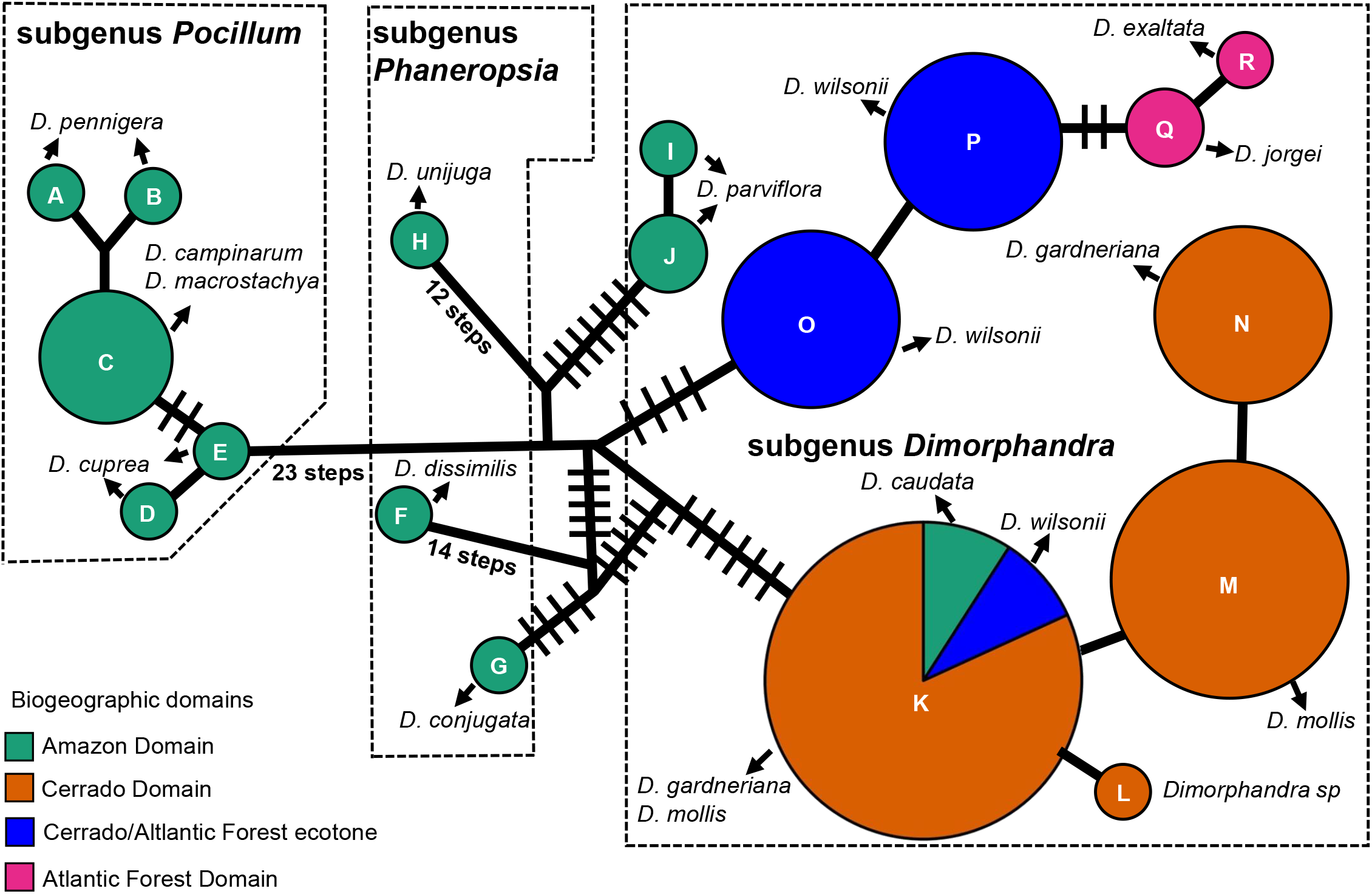
Median-joining network for 56 specimens of *Dimorphandra*. The dataset (446 pb) was resulted from nuclear region (ITS). Each circle represents a given haplotype (coded with number); size is proportional to relative frequencies. Numbers of mutational steps are indicated with bars when there is more than one (unless indicated otherwise)

In agreement with the cpDNA network, the ITS network also showed that the haplotypes recovered from the Amazonian species of *Dimorphandra* (11 haplotypes) required a larger number of mutational steps to connect to the remaining of the network.

Meanwhile, connections of the haplotypes of the Cerrado, the Cerrado/Atlantic Forest ecotone, and the Atlantic Forest (8 haplotypes) required few mutational steps (up to 2). There was a haplogroup that brought together four haplotypes: the central haplotypes (O and P) were both found in *D. wilsonii* (a species from the Cerrado/Atlantic Forest ecotone), the tip haplotypes (Q and R) were found in *D. jorgei* and *D. exaltata*, respectively (both species are from the Atlantic Forest). The second haplogroup also contained four haplogroups (K, L, M and N), all of which were from the Cerrado species (*D. gardneriana* and *D. mollis*). There was only one haplotype sharing among domains. One Amazonian species (*D. caudata*) shared haplotype K with two species of the Cerrado (*D. gardneriana* and *D. mollis*) and the species from the Cerrado/Atlantic Forest ecotone (*D. wilsonii*).

### Ancestral area inferences

The BBM analysis reconstructed the ancestral area for the 17 species of *Dimorphandra* (Fig. 5b). The Amazon was the most likely ancestral area for species of two subgenera: *Phaneropsia* (*D. conjugata*, *D. dissimilis*, and *D. unijuga*) and *Pocillum* (*D. campinarum*, *D. coccinea*, *D. cuprea*, *D. macrostachya*, *D. pennigera*, *D. polyandra*, and *D. vernicosa*). Also, the Amazon was the ancestral area of *D. parviflora*, a species that belongs to the subgenus *Dimorphandra*. Interestingly, within the subgenus *Dimorphandra*, the Amazon was the most likely ancestral area for three species. Two of them occurred in the Cerrado (*D. mollis* and *D. gardneriana*) and the third species in the Cerrado/Atlantic Forest ecotone (*D. wilsonii*). Most likely, the ecotone was the ancestral area of the two species (*D. jorgei* and *D. exaltata*) that occurred in the Atlantic Forest.

**Fig.5.**
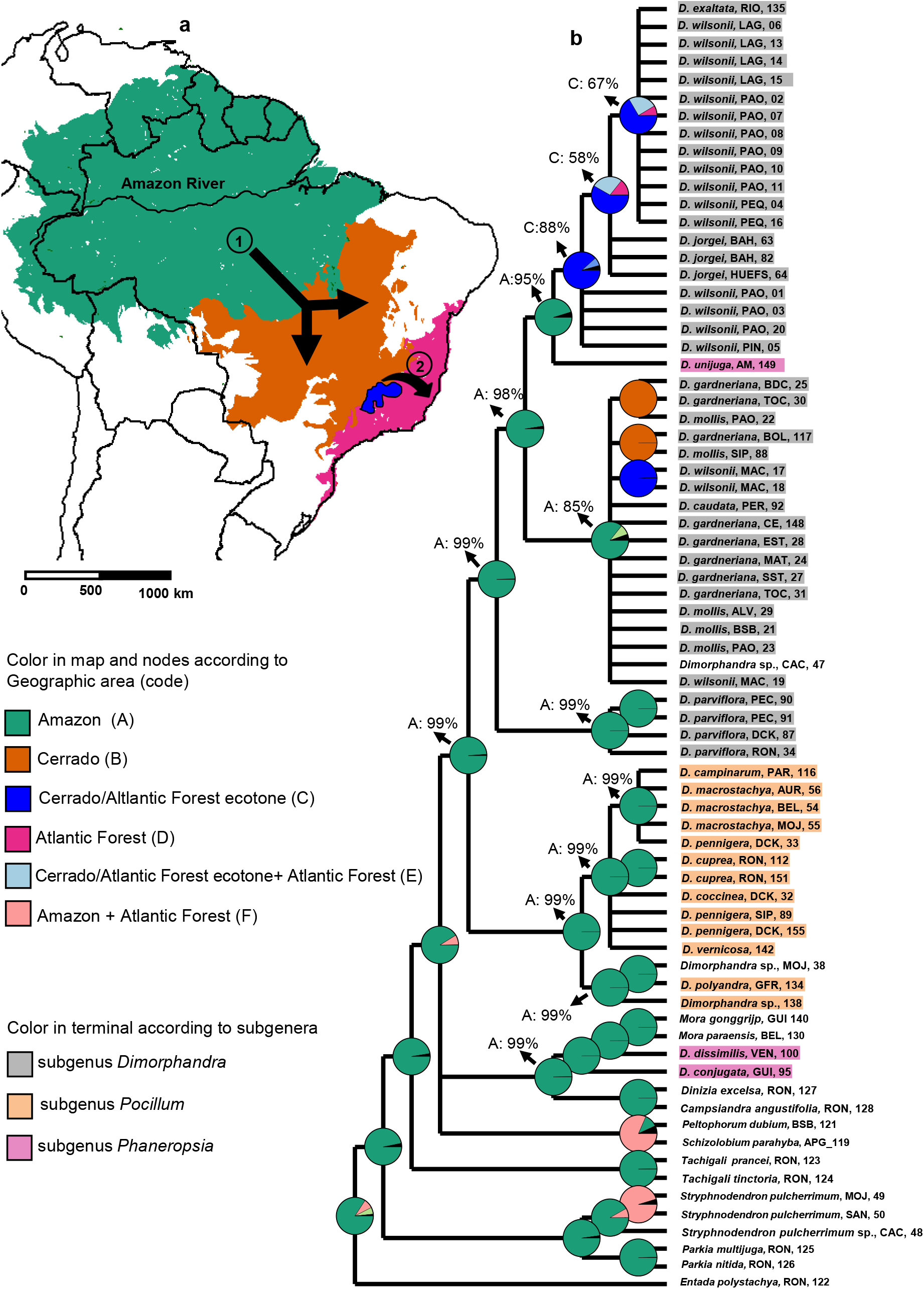
Ancestral area reconstruction for genus *Dimorphandra*. **a** The map shows the current distribution range of the genus *Dimorphandra* in South America, and arrows indicate major dispersal events inferred in this study: (1) from the Amazon into the Cerrado; (2) from the Cerrado into the Atlantic Forest. **b** Ancestral area reconstruction obtained from Bayesian Binary MCMC (BBM) analysis. Pie charts at each node represent the probabilities of geographic distribution of the hypothetical ancestor. Arrows indicated the most probable ancestral area and probability values inmajor nodes.

### Ecological Niche Modeling

To investigate the habitat suitability for *Dimorphandra* within the three domains (Amazon, Cerrado, and Atlantic Forest), we carried out projections for two species from the Amazon (*D. macrostachya* and *D. pennigera*), two species from the Cerrado (*D. mollis* and *D. gardneriana*), one species from ecotone Cerrado/Atlantic Forest (*D. wilsonii*), and two species from the Atlantic Forest (*D. exaltata* and *D. jorgei*). The AUC, TSS and COR values indicated that the generated models performed well to predict species occurrence in relation to the bioclimatic variables (Supplementary Table S6).

The projections (Fig. 6) covered the following time periods: Last Interglacial (LIG, 130 kyr BP), Last Glacial Maximum (LGM, 21 kyr BP) and Present. Projections with either *D. macrostachya* (Fig. 6a) or *D. pennigera* (Fig. 6b) showed the presence of areas highly suitable in northwestern South America during LIG; these areas extended northeastward during LGM and were maintained at the present. During LIG and LGM, there were no areas with high suitability for *D. macrostachya* (Fig. 6a) and *D. pennigera* (Fig. 6b) in the Cerrado and Atlantic Forest.

**Fig.6.**
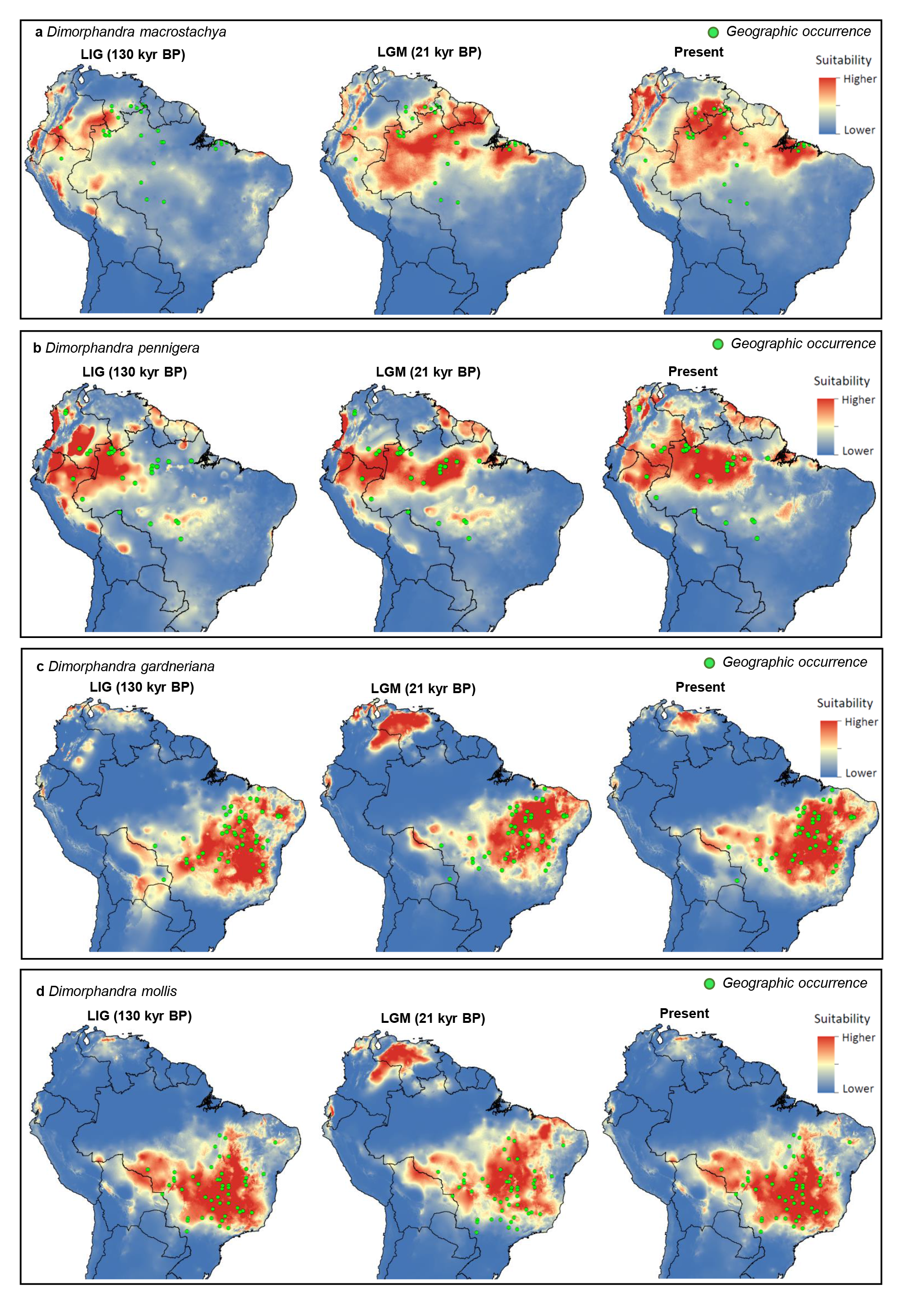
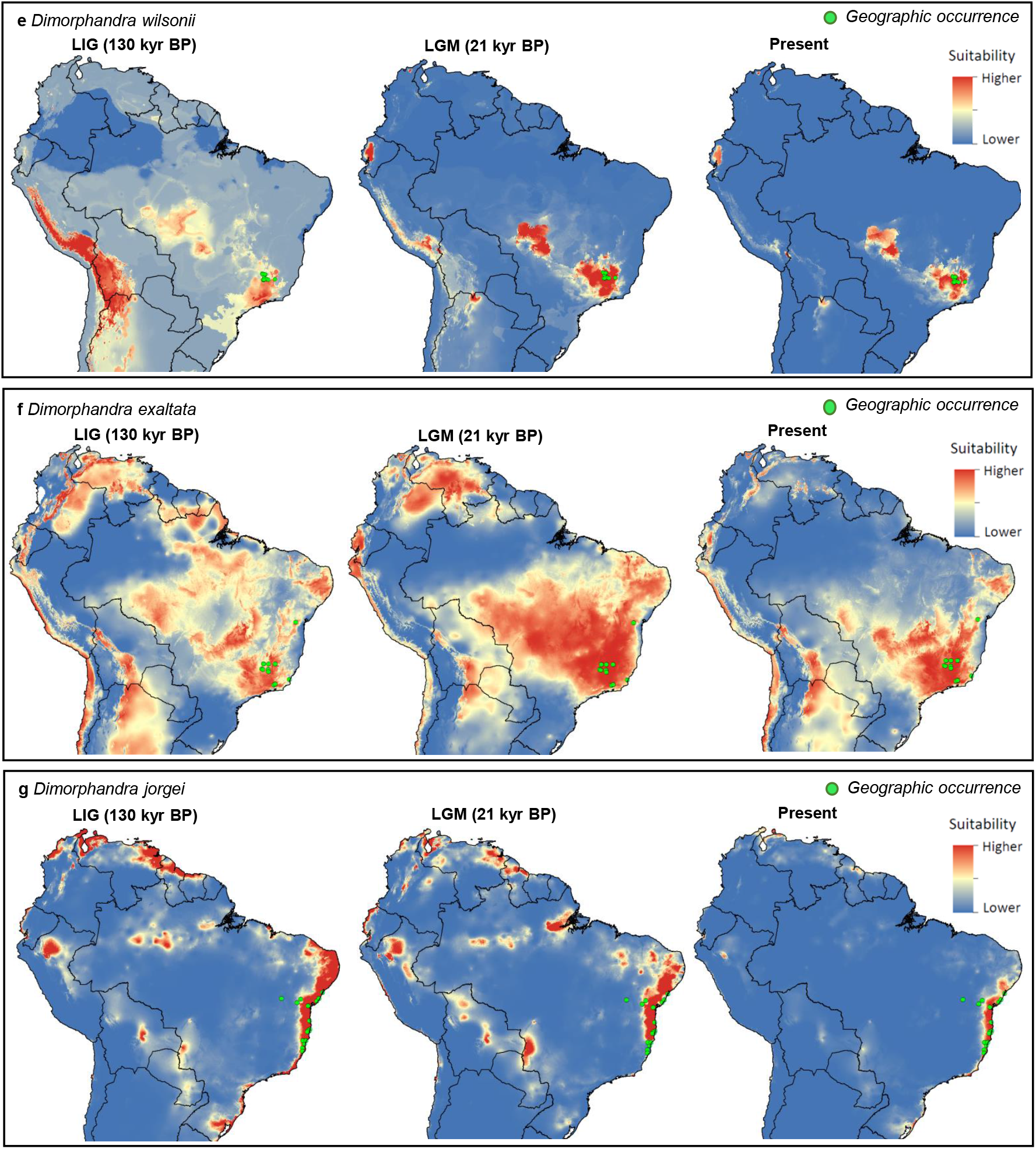
Ecological Niche Modeling for seven species of *Dimorphandra* in South America during the Last Interglacial (LIG, 130kya BP), the Last Glacial Maximum (LGM, 21 kya BP), and the Present. **a** *D. macrostachya*. **b** *D.pennigera*. **c** *D. gardneriana*. **d** *D. mollis*. **e** *D. wilsonii*, **f** *D. exaltata*, **g** *D. jorgei*.

Meanwhile, projections for *D. mollis* (Fig. 6c) and *D. gardeneriana* (Fig. 6d) revealed high habitat suitability in central Brazil (Cerrado) overtime (LIG, LGM and present); but during LGM there was a large area potentially suitable in northern South America, which coincides with Llanos, Savana from Colombia, and Venezuela.

Projections for *D. wilsonii* (Fig. 6e), *D exaltata* (Fig. 6f) and *D. jorgei* (Fig. 6g) indicated the existence of fragmented areas with moderate-high suitability spread across the South America during LIG and LGM.

## DISCUSSION

### Incomplete lineage sorting of ancient polymorphisms

As it is presently circumscribed (Silva 1986), the genus *Dimorphandra* is subdivided into three subgenera: (1) *Dimorphandra*, with 11 species, (2) *Phaneropsia*, with five species, and (3) *Pocillum*, with 10 species. Phylogenetic placement of species that belong to the subgenus *Phaneropsia* was incongruent between phylogenic reconstructions that used data from either the cpDNA (Fig. 1) or the nuclear genome (the ITS gene region) (Fig. 2).

Our phylogenic reconstruction based on cpDNA data suggested that the cpDNA genomes of two species (*D. conjugata* and *D. dissimilis*) that are currently assigned to the subgenus *Phaneropsia* were more closely related to species of the genus *Mora* (*M. paraensis* and *M. gonggrijpii*) than to the remaining congeners. This relationship was also recovered in previous phylogenetic studies, where a single species from subgenus *Phaneropsia* (*D. conjugata*) grouped within a clade containing members of the legume genera *Burkea*, *Mora* and *Stachyothyrsus* (Bruneau et al. 2008; Manzanilla and Bruneau 2012). However, the reduced sampling of Dimorphandra in these studies did not allow testing its monophyly.

A certain degree of close proximity between *Mora* and the subgenus *Phaneropsia* has been proposed previously (Bentham and Hooker 1867). There were several suggestions either to include *Mora* together with the subgenus *Phaneropsia* (Taubert 1864; Benthan and Hooer 1867) or to place *Mora* together with the subgenus *Phaneropsia* into a novel taxonomic entity (Ducke 1925; Sandwith 1932; Amshoff 1939; Silva 1986). In sharp contrast with the uncertain placement of the subgenus *Phaneropsia*, the other two subgenera (*Dimorphandra* and *Pocilum*) have always been considered as distinct entities, with well delimited morphologically based taxonomic assignments (Silva 1986).

Another incongruent placement took place with *D. unijuga* (subgenus *Phaneropsi*a), as it came together with species of the subgenus *Dimorphandra* in the cpDNA tree (Subclade IIIC; Fig 1). Morphologically, both leaflet shape and inflorescences of *D. unijuga* exhibited a certain affinity with those of *D. jorgei* (subgenus *Phaneropsia*). However, fruits and leaves clearly distinguish *D. unijuga* apart from *D. jorgei* (Silva 1986). This relationship was not recovered in the ITS tree (Fig 2).

At least two plausible, non-mutually exclusive scenarios may explain the incongruent placements we detected in the subgenus *Phaneropsia*: hybridization events that led to chloroplast capture; and incomplete lineage sorting.

Chloroplast capture is a common cause for inconsistencies between phylogenetic reconstructions based on nuclear and cpDNA marker sequences (Rieseberg and Soltis 1991). Chloroplast captures may be associated with geographic locations more than taxonomic relationships (Rieseberg and Soltis 1991). To explain the patterns, we identify in the subgenus *Phaneropsi*a, chloroplast capture in Clade I had to occur independently from that in Subclade IIIC (Fig. 1). Moreover, the four specimens of Clade I (GUI 140, *M. paraensis;* BEL 130, *M. gonggrijpii;* VEN 100, *D. dissimilis*; and GUI 95, *D. conjugata*) were sampled from relatively highly distant geographic locations across the Amazon (Supplementary Table S1). The Amazonian *D. unijuga* detained highly differentiated cpDNA markers relative to the Atlantic species *D. wilsonii* and *D. jorgei*. Thus, such findings suggested chloroplast capture as a less probable scenario to explain fully the conflicting gene trees we uncovered.

Incomplete lineage sorting of ancient polymorphisms, however, may be a more plausible cause for the incongruence we detected in *Dimorphandra*. According to this scenario, the genus *Mora* and species of the subgenus *Phaneropsi*a inherited their chloroplast genomes from a common ancestor that existed before the diversification events that gave rise to both *Mora* and *Dimorphandra*. This scenario implies that the subgenus *Phaneropsi*a detains a basal position within *Dimorphandra*; thus explaining the close proximity between subgenus *Phaneropsi*a and the genus *Mora*. Incomplete lineage sorting has been documented in several Leguminosae lineages, such as *Lespedeza* (Xu et al. 2012) and *Acacia acuminata* (Byrne et al. 2002). The incongruent placement of *D. unijuga* seems intricate and difficult to explain while assuming a single scenario of either hybridization or lineage sorting.

Our analyses, based on ITS sequences, suggested that *Dimorphandra* may be considered as a monophyletic genus. However, a reassessment based on a large sample of the nuclear genome (e.g. Koenen et al. 2020) is needed to confirm its monophyly. This new phylogenomic assessment will require a dense taxonomic sampling, including species from both *Mora* and *Dimorphandra*, especially species from the subgenus *Phaneropsia*.

### An *Amazonian origin for Dimorphandra*

The deep evolutionary lineages we detected in the Amazon provided strong evidence for the extant lineages of *Dimorphandra* have an origin in that biogeographic domain. Indeed, the Amazon is the domain that contained the highest diversity of extant species of *Dimorphandra* (Silva 1986; Silva 2019). The Amazon has been proposed to the likely origin of many other plant lineages (e.g. Schley et al. 2018), including taxa closely related to *Dimorphandra*, such as *Parkia* (Conceição Oliveira et al. 2021).

Habitat shifts were shown to have played important roles in shaping the diversity of both flora and fauna across the three main biogeographic domains of South America: Amazon, Cerrado, and Atlantic Forest (Simon and Pennington 2012; Terra-Araujo et al. 2015; Simon et al. 2016; Souza-Neto et al. 2016; Antonelli et al. 2018; Aguiar et al. 2019). Our results suggested that habitat shifts also contributed to the diversification of *Dimorphandra*. Two subgenera (*Phaneropsia* and *Pocillum*) retained their ancestral, forested habitat in the Amazon; whereas the subgenus *Dimorphandra* took part in a dynamic process of habitat shift, from forests to savannas, which seem to have occurred relatively recently in the evolutionary time.

### Habitat shift and the diversification of Dimorphandra

Several studies postulated past climate changes as driving forces that shaped the Neotropics through recurring cycles of alternating cool–dry and warm–wet climates, which in turn promoted episodes of expansion or contraction of vegetations (Behling and Hooghiemstra 2001; Bueno et al. 2017). Recurrent cycles were proposed to have occurred during the last 2 million years of the Pleistocene (Whitmore and Prance 1987), but they also may have occurred much earlier (Mayle 2004; Pennington et al. 2004). During more humid episodes, therefore, *Dimorphandra* from the Amazon may have dispersed towards the neighboring areas of the Cerrado. Whilst during drier episodes; *Dimorphandra* from the Cerrado may have entered areas that closed-canopy vegetation, such as the Amazon and Atlantic Forest, occupied previously. When considered over evolutionary timescales, the recurrent range shifts certainly had the potential to trigger adaptation and provided opportunities for speciation to take place in *Dimorphandra*. Moreover, ecotones that form during such alternating cycles owing to climate changes may have been dynamically active over evolutionary timescale; temporary corridors may have connected the Atlantic coast to the Amazon through the Cerrado of Central Brazil (Auler et al. 2004; Mayle 2004; Carnaval and Moritz 2008). Ancient connections across the three domains (Amazon, Cerrado, and Atlantic Forest) have been proposed to have occurred recurrently, from the Miocene to the Pleistocene (Oliveira-Filho and Ratter 1995; Ledo and Colli 2017; Machado et al. 2018). Over time, important migration routes could allow for the dispersal of many animal and plant species (Buzatti et al. 2018; Fabrício Machado et al. 2021), including *Dimorphandra*.

The Amazon seems to be the likely source of the lineage that gave rise to the savanna species of *Dimorphandra*. This intriguing scenario requires an Amazonian lineage to undergo habitat shift from closed-canopy vegetation, such as the Amazonian forests, to the open vegetation (savanna), such as the Cerrado. Over evolutionary time scales, such parental lineage underwent speciation events and originated the extant species that occur in the Cerrado: *D. mollis*, *D. gardneriana*, and *D. wilsonii*. The sister relationship (Fig. 1 and Fig. 2) between an Amazonian species (*Dimorphandra parviflora*) and the clade of extra-Amazonian congeners may be regarded as a reminiscent clue of the ancient phylogenetic link that connected a putative ancestral lineage of Amazonian origin and the more recently derived lineages that dispersed to the East and today inhabit the Cerrado and the Atlantic Forest.

In addition to *Dimorphandra*, many other plant lineages likely underwent habitat shifts, from the Amazon to the Cerrado. Those plant taxa include *Andira* and *Eriotheca* (Simon and Pennington 2012), *Parkia* (Conceição Oliveira et al. 2021), *Stryphnodendrom* (Simon et al. 2016). Animal taxa may also have experienced habitat shifts, such as *Phyllomus* (Machado et al. 2018). Thus, our findings add to the growing evidence that many plant lineages of the Cerrado detain an Amazonian origin (Simon and Pennington 2012; Antonelli et al. 2018).

*Dimorphandra* and other plant lineages – such as *Myrcia* sect. *Aguava* (Lima et al. 2021) – seem to have shifted habitats from the Cerrado to the Atlantic Forest. Haplotype placements in the networks (Fig. 3 and Fig. 4) together with species arrangements in the phylogenetic reconstructions (Fig. 1 and Fig. 2) allow us to postulate that the extant species *D. exaltata* and *D. jorgei* likely originated from a parental lineage from either the adjacent Cerrado or the ecotone Cerrado/Mata Atlântica.

The origin of *D. caudata* is highly intriguing. In our phylogenetic reconstructions, *D. caudata* seem an out-of-place species. The species occur in the Amazon, but is phylogenetically more closely related to congeners of the Cerrado (*D. mollis* and *D. gardneriana*) and the ecotone Cerrado/Atlantic Forest (*D. wilsonii*). One possible scenario for the origin of *D. caudata* requires an ancestor species in the Cerrado; thus, explaining the close phylogenetic relationship between *D. caudata* and the extant Cerrado congeners. According to this scenario, the ancestor of *D. caudata* underwent a habitat shift from the Cerrado back to the Amazon, followed an adaptation to the close-canopy Amazonian forests. Transitions from an ancestral area in the Cerrado followed by multiple colonization events to the Amazon and Atlantic Forest have been documented for Vochysiaceae (Gonçalves et al. 2020).

### Extra-Amazonian species of Dimorphandra are genetically depauperate

In contrast to the highly polymorphic and divergent lineages we uncovered in the Amazon, the lineages from either the Cerrado or the Atlantic Forest were genetically depauperate. Previous studies also pointed out low levels of genetic diversity in *D. exaltata*, *D. mollis*, and *D. wilsonii* (Viana e Souza and Lovato 2010; Vinson et al. 2015; Muniz et al. 2019). In our phylogenetic reconstructions (Figs. 1 and 2), the extra-Amazonian species from the Cerrado (*D. mollis* and *D. gardneriana*), ecotone (*D. wilsonii*), and Atlantic Forest (*D. exaltata* and *D. jorgei*) were placed within derived clades, all of which with very short internal branches. Additionally, network analyses (Fig. 3 and Fig. 4) indicated that the extra-Amazonian species detained low levels of haplotype differentiation and haplotype diversity; connection among those haplotypes occurred through few (1 to 3) mutational steps. In consonance with our finding that the extra-Amazonian species are genetically depauperate and seem to detain low levels of genetic differentiation. Taken together, those results suggested a recent origin for the extra-Amazonian lineages of *Dimorphandra*, with rapid diversification taking place afterwards. This scenario can explain the relative low levels of genetic diversity we detected in the extra-Amazonian species and support a previous suggestion that they exhibit great morphological overlap (Silva1986).

### A biogeographical hypothesis for the dispersion of the subgenus Dimorphandra

Based on our findings, we propose two major dispersal events that may account for the origin and current geographic distribution of the subgenus *Dimorphandra* (Fig. 5a). Firstly, an Amazonian ancestor lineage dispersed into the Cerrado and afterwards gave rise to the current congeners *D. mollis, D. gardneriana*. It is not yet clear whether this dispersal took place through a single colonization wave; i.e., an ancestral population migrated from the Amazon into the Cerrado and later gave rise to the two extant haplogroups uncovered (Fig. 3). Or whether there was more than one colonization wave; i.e., at least two ancestral populations migrated from the Amazon to the Cerrado at different time periods; each of which giving rise to one of the two extant haplogroup (Fig. 3). A second dispersal event took place from the Cerrado towards the Atlantic Forest, giving rise to the extant species *D. exaltata* and *D. jorgei* (Fig. 5a). According to this scenario, those two species are not evolutionary relicts, but rather they are recently derived species; their narrow range resulted from their recent origin and specialization to the closed-canopy vegetation of the Atlantic forests.

### Implications for conservation

Our study lays a foundation for understanding the evolutionary history of the genus *Dimorphandra*. The five extra-Amazonian species (*D. gardneriana*, *D. mollis*, *D. wilsonii*, *D. exaltata*, and *D. jorgei*) merit attention of conservation programs. Currently, the natural habitats of those species have been reduced significantly due to either urban development or land conversion to agriculture across their range. Naturally rare, *D. wilsonii* is listed as critically endangered according to National Center for Plant Conservation (CNCFlora). Both *Dimorphandra exaltata* and *D. jorgei* are also considered rare species (Silva 1986), however their complete status have not been fully addressed. On other hand, *D. gardneriana* and *D. mollis* showed a widespread distribution (Silva 1986); their fruits have been widely used as a source of the flavonoid rutin, an agent of pharmacological interest (Cunha et al 2009; Viana e Souza and Lovato 2010). The unmanaged exploitation of those natural resources may lead to a decrease in the genetic diversity of the species (Gonçalves et al. 2010).

So far, *D. wilsonii* is the only congener that has been included in a national plan for conservation (Fernandes et al. 2014). The conservation programs may also consider including *D. gardneriana*, *D. mollis*, *D. exaltata*, and *D. jorgei*. The establishment of ecological corridors is apparently a suitable strategy to connect populations of *Dimorphandra* that are isolated across a fragmented landscape. Thus, such corridors could facilitate the translocation of dispersers and pollinators, increasing gene flow and genetic diversity in those species of *Dimophandra*.

## Acknowledgements

We thank the many herbaria and researchers that provided us with samples for DNA extraction: Jardim Botânico do Rio de Janeiro (RB), Missouri Botanical Garden (MO), Herbário da Amazônia Meridional (HERBAM), Herbário da Guiana (CAY), Smithsonian Institution (US), Herbário da Embrapa Cenargen (CEN), Herbário da Universidade Estadual de Feira de Santana (HUEFS), Herbário da Universidade Federal de Viçosa (VIC), and Herbário da Embrapa Amazônia Oriental (IAN), as well as Regina Celia Martins da Silva, Karen Redden, Luciano Margalho, Claúdia Pombo, Chieno Suemitsu, Silvana Monteiro and Lilian Procópio. This work was supported by grants of the Minas Gerais State Foundation of Research Aid – FAPEMIG (APQ-00150-17) and by The National Council of Scientific and Technological Development – CNPq to LOO (PQ 302336/2019-2) and MFS (PQ305570/2021-8). VDR received a scholarship from CNPq (130903/2018-3).

## Material Supplementary

**Fig.S1.** Maximum likelihood phylogeny (consensus tree) showing relationship among 17 species of *Dimorphandra* (59 specimens) and nine related genera (14 specimens). The dataset (2615 pb) was obtained from the concatenation of five cpDNA regions: an intron (trnL-UAA), three intergenic spacers (*trn*L–*trn*F; *trn*H–*psb*A and *psb*B–*psb*F) and a region of the gene Maturase K. Branch lengths are drawn to scale. Nodal support values are given as bootstrap values when ≥ 85. Scale bar corresponds to the expected number of substitutions per site

**Fig.S2.** Maximum likelihood phylogeny (consensus tree) showing relationship among 14 species of *Dimorphandra* (44 specimens) and seven related genera (9 specimens). The dataset (638 pb) resulted from nuclear region (ITS). Branch lengths are drawn to scale. Nodal support values are given as bootstrap values when ≥ 85. Scale bar corresponds to the expected number of substitutions per site

