## Supplementary figures and images for "FROM FOREST TO SAVANNA AND BACK TO FOREST: EVOLUTIONARY HISTORY OF THE GENUS *Dimorphandra* (LEGUMINOSAE)"

### Supplementary Figures

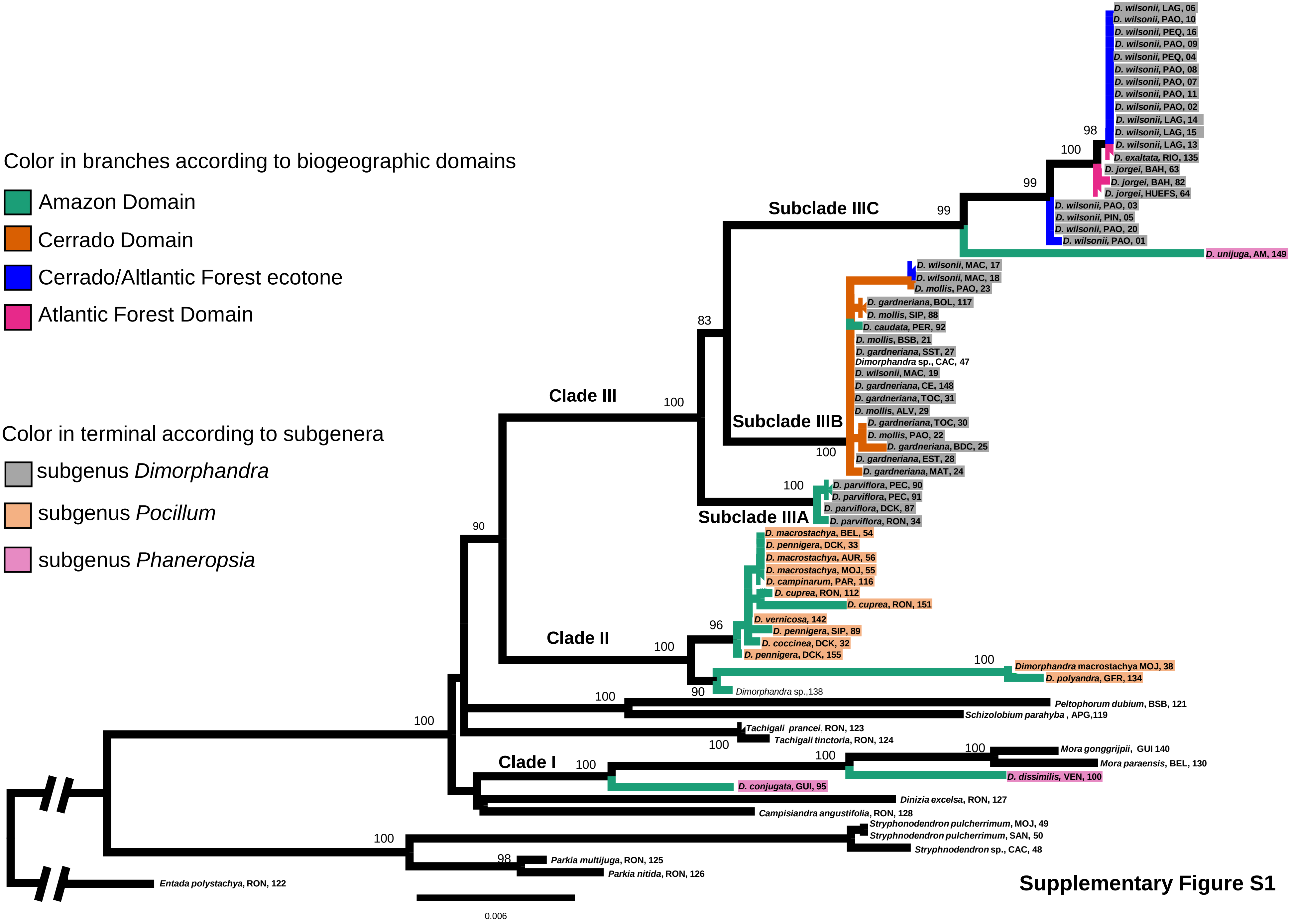


**Supplementary Figure S1**

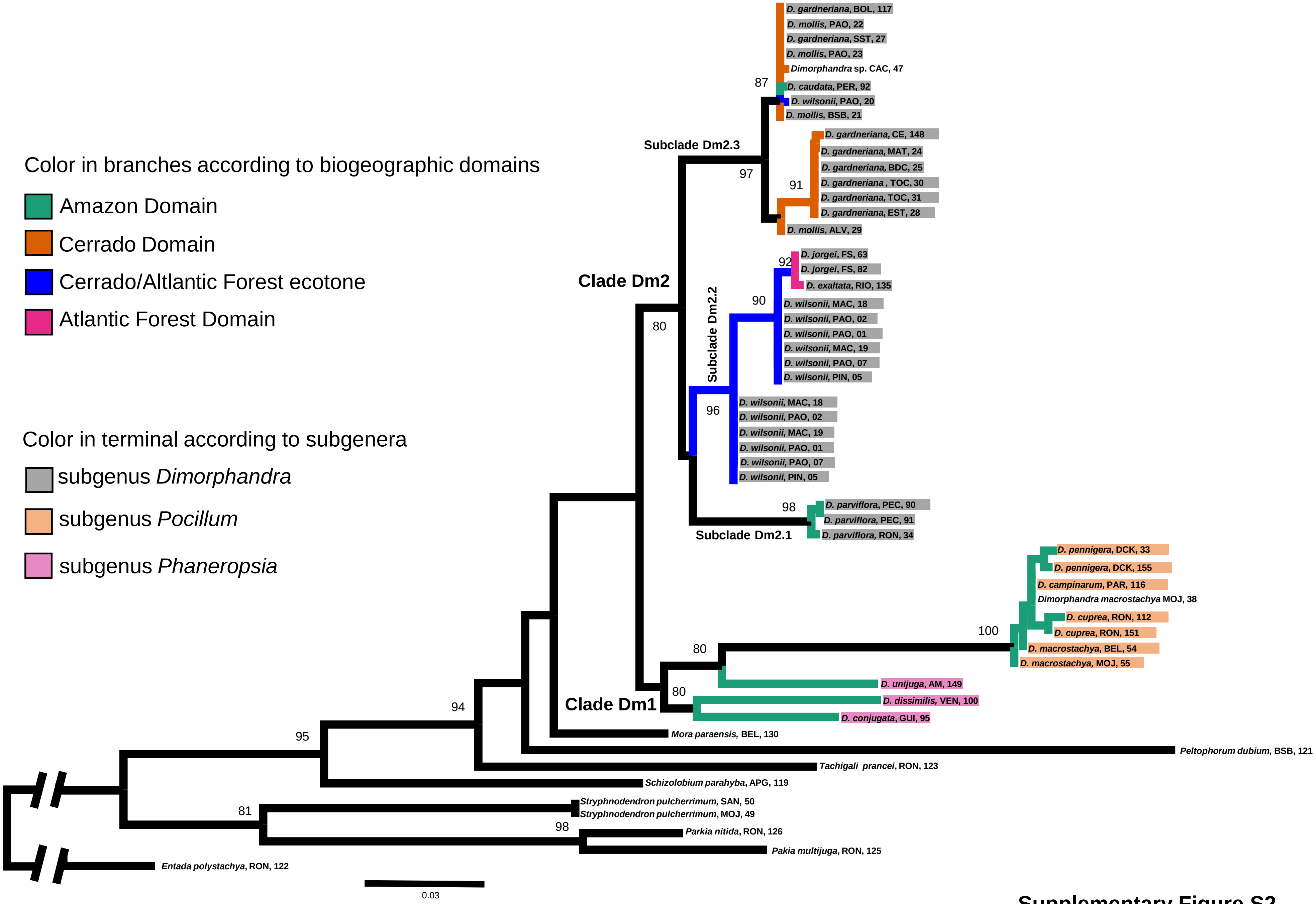


**Supplementary Figure S2**
